# Interferon-□ Exposure of Human iPSC-derived Neurons Alters Major Histocompatibility Complex I and Synapsin I Protein Expression

**DOI:** 10.1101/2021.12.15.472810

**Authors:** Adam Pavlinek, Rugile Matulevicute, Laura Sichlinger, Lucia Dutan Polit, Nikolaos Armeniakos, Anthony C. Vernon, Deepak P. Srivastava

## Abstract

Human epidemiological data links maternal immune activation during gestation with increased risk for neurodevelopmental disorders including schizophrenia. Animal models of maternal immune activation (MIA) provide causal evidence for this association and strongly suggest that inflammatory cytokines act is a critical link between maternal infection and aberrant offspring brain and behavior development. This includes evidence for reduced synapse formation, consistent with *post-mortem* and *in vivo* evidence of reduced synaptic density in schizophrenia. However, to what extent specific cytokines are necessary and sufficient for these effects remains unclear. Using a human cellular model, we recently demonstrated that acute exposure to interferon-□ (IFN□) recapitulates molecular and cellular phenotypes associated with neurodevelopmental disorders. Here, we extend this work to test whether IFN□ affects synapse formation in an induced neuron model that generates forebrain glutamatergic neurons. Using immunocytochemistry and quantitative PCR, we demonstrate that acute IFN□ exposure results in significantly increased MHCI expression at the mRNA and protein level. Furthermore, acute IFN□ exposure decreases synapsin I protein in neurons but does not affect synaptic gene mRNA levels. Interestingly, complement component 4A (*C4A*) mRNA is also significantly increased following acute IFN□ exposure. This study builds on our previous work by showing that IFN□-mediated disruption of relevant synaptic proteins can occur at early stages of synapse formation, potentially contributing to neurodevelopmental disorder phenotypes such as schizophrenia.

## Introduction

Human epidemiological studies and animal models suggest a link between maternal immune activation (MIA) and an increased risk for psychiatric disorders with a putative neurodevelopmental origin, including schizophrenia (Kępińska et al., 2020). Although there are likely to be many plausible factors that are critical for establishing neurodevelopmental resilience or susceptibility to MIA (Meyer, 2019), there is evidence to suggest that the intensity of the maternal immune response is an important factor linking maternal infection to abnormal brain development and behavioural phenotypes (Mueller et al., 2018;Mueller et al., 2019;Estes et al., 2020). MIA models present with deficits in cognitive and social behaviours (Mueller et al., 2021), which are accompanied by altered synaptic plasticity, decreased synaptic protein levels, and reduced dendritic spine density, predominantly in the prefrontal cortex and hippocampus (Coiro et al., 2015;Weir et al., 2015;Zhang and van Praag, 2015;Giovanoli et al., 2016;Pekala et al., 2021). These findings are consistent with *in vivo* neuroimaging evidence for reduced synaptic density, as measured by SV2A-PET in schizophrenia (Onwordi et al., 2020), reduced dendritic spines (Glantz and Lewis, 2000), and decreased expression of synaptic proteins in *post-mortem* brain tissue from individuals with schizophrenia (Osimo et al., 2019).

One key feature of the maternal immune response that shapes these phenotypes is the elevation of numerous cytokines in the maternal serum, placenta and foetal brain (Urakubo et al., 2001;Garay et al., 2013). Consistent with this view, elevated cytokine levels in maternal serum are predictive for the risk of offspring developing schizophrenia (Allswede et al., 2020). Moreover, human birth cohort studies provide evidence that variation in maternal cytokine levels during pregnancy are associated with offspring cognitive and behavioral outcomes, as well as measures of fetal brain growth and connectivity as measured by MRI (Graham et al., 2018;Rudolph et al., 2018;Rasmussen et al., 2021). Collectively, these studies suggest that changes in maternal cytokines during pregnancy can have long-lasting consequences. To what extent specific cytokines are necessary and sufficient for these effects remains unclear, and the underlying molecular mechanisms remain to be fully elucidated. Evidence from animal models of MIA provides strong support for the involvement of interleukins, particularly IL-6, but also TNF-alpha, IL-1beta, IL-10 and interferon-□ (IFN□) (Smith et al., 2007;Lins et al., 2018;Mirabella et al., 2021;Mueller et al., 2021). IFN□ has been found to have increased levels in the plasma of first-episode schizophrenia patients (Lesh et al., 2018). However, it is unclear whether and how this is contributing to increased risk for schizophrenia. In addition to its key role in the response to viral infection, IFN□ has also been shown to induce retraction of dendrites and inhibit synapse formation in the central nervous system (Kim et al., 2002;Monteiro et al., 2017).

We previously showed that exposure of neural progenitor cells (NPCs) and developing immature neurons derived from human induced pluripotent stem cells (iPSCs) to IFN□ results in gene expression changes in genes associated with schizophrenia and autism, and altered neuronal morphology, with increased outgrowth in exposed neurons (Warre-Cornish et al., 2020). In particular, IFN□ treatment increases major histocompatibility complex I (MHCI) expression (Warre-Cornish et al., 2020). Class I MHC family molecules are best known for their function in presenting antigens to T-cells (Shatz, 2009). MHCI is expressed in neurons and neural progenitors and has been found to be important in neuronal plasticity and for the co-regulation of synapse pruning in mice (Shatz, 2009;Lee et al., 2014). MHCI negatively regulates synapse density in developing cortical neurons, with *in vitro* manipulations of MHCI expression inversely affecting the density of both GABAergic and glutamatergic synapses in rat and mouse cultures (Glynn et al., 2011). In a mouse model of MIA, synapse number in cultured cortical neurons was decreased, and MHCI was found to be required for this MIA-induced effect on synapse density (Elmer et al., 2013). However, it is not known if cytokine-driven increases in MHCI impact synapse formation in human neurons. Genome-wide association studies also support the MHC loci links with schizophrenia (Shi et al., 2009;Ripke et al., 2014). For example, variation of complement component 4 (C4) at the MHCIII locus and human leukocyte antigen-B (HLA-B) at the MHCI locus is strongly associated with increased risk for schizophrenia (Sekar et al., 2016).

In our previous work, gene expression changes following IFN□ treatment included increased expression of MHCI genes and downregulation of synapse-related genes in exposed iPSC-derived immature neurons (Warre-Cornish et al., 2020). Given that IFN□ has been shown to affect expression of synaptic genes in iPSC-neurons in the absence of glial cells (Warre-Cornish et al., 2020), we wanted to determine whether the Neurogenin 2 optimized inducible overexpression ioGlutamatergic cell line (Pawlowski et al., 2017) could be a useful system for studying the effects of IFN□ in a pure population of glutamatergic cells. Using this system, we aim to further characterize the effect of IFN□ treatment on MHCI expression and synapses. We find that IFN□ exposure increases MHCI protein staining and *HLA-B* and *C4A* expression but decreases the synaptic protein synapsin I in cell bodies without altering the expression of synaptic genes.

## 1 Methods

### 1.1 HiPSC culture, neuralization, and treatment

The ioGlutamatergic male neurotypical stem cell line (Pawlowski et al., 2017) was obtained from BitBio (Cambridge, UK) under MTA agreement. ioGlutamatergic cells were maintained in Stemflex media (Gibco; A3349401) on six-well plates coated with 1:100 Geltrex (Life technologies; A1413302). Media was changed every 48 hours and passaged when 70-80% confluent with HBSS and Versene (Gibco; 15040066) at 37°C before being transferred into new Stemflex medium. Neuralization was conducted based on the protocol used by Pawlowski et al. (2017). Cells for experiments were terminally plated onto 6-well plates (for RNA extraction) or glass coverslips in 24-well plates (for immunocytochemistry) coated with Poly-D-Lysine (5 μg/ml, PDL, A-003-E; Millipore) and laminin (1 mg/ml Sigma L2020). iPSCs were dissociated with accutase (A11105-01; Thermo Fisher Scientific) before being diluted with medium and subsequently resuspended in N2 medium (supplementary table 1) with 1μg/ml doxycycline hyclate and 10 μM ROCK inhibitor (Sigma; Y0503). Cells were plated at a density of 900,000 cells/well for RNA extraction and 25,000 cells/well for ICC. The cells were incubated at 37**°**C; 5% CO2; 20% O2 with daily N2 media changes supplemented with 1μg/ml doxycycline hyclate. 25 ng/ml IFN□ (Abcam, AB9659; diluted in DMEM) for treatment conditions or vehicle (DMEM) was added at day 3 to the N2 medium. The cells were incubated for 24-hours before sample collection (Warre-Cornish et al., 2020).

The 127_CTM_01 iPSC male neurotypical line (Adhya et al., 2021) was differentiated into NPCs using a dual SMAD inhibition protocol (Shum et al., 2020;Adhya et al., 2021). Briefly, the NPCs were expanded from day 18 frozen stocks in maintenance medium (1:1 N2:B27, 10ng/ml bFGF) for seven days. Before treatments the cells were plated on 12-well NUNC™ tissue culture plates (Thermo Scientific; 150628) at a density of 500,000 cells/well, with dedicated wells for treatment and vehicle treatments. The day after plating, the cells were exposed to 25 ng/ml IFNγ or vehicle and incubated for 24-hours before sample collection.

### 1.2 Immunocytochemistry

Cells were fixed with 4% formaldehyde in PBS-sucrose for 10 minutes at room temperature, washed 2× with Dulbecco’s PBS (DPBS, Gibco), and then fi xed with ice cold Methanol at 4°C for 10 minutes, then washed 2× with DPBS. Cells were permeabilized and blocked using 2% normalized goat serum (NGS) in DPBS with 0.1% triton x-100 for 2 hours. Antibody solutions (supplementary table 2) were prepared in 2% NGS in DPBS. The coverslips were incubated with primary antibody solution at 4°C overnight, then washed 3× with DPBS for 10 minutes each and incubated with secondary antibodies for 1 hour at room temperature. The coverslips were washed 3× with DPBS for 10 minutes each and incubated for 5 minutes in DAPI solution, followed by two DPBS washes, then mounted onto glass slides using ProLong Gold antifade reagent (Invitrogen P36930).

### 1.3 Microscopy and image analysis

Coverslips were imaged using a Leica SP5 confocal microscope. The gain and other imaging parameters were set using the vehicle control and were not changed during subsequent imaging of the control and IFN□ exposed coverslips. 246.5×246.5 μm regions were imaged. The Z stack thickness was kept at 0.5 μm. Z stacks were maximally projected to form a single image in FIJI. Prior to measuring fluorescent intensity, the background of each image and channel was measured in FIJI by selection of 10 25×25-pixel areas of background and by then measuring the mean intensity and standard deviation (SD) of intensity of each area. The mean of these measurements + 2SD was then subtracted from the image. Cell Profiler (Carpenter et al., 2006) was used to identify the nuclei, cells, cell bodies, processes, and the cytoplasm and to measure the mean intensity of the MHC and synapsin I channels. Mean intensity values of 0 were excluded from the analysis. The pipeline is provided as a supplementary file.

### 1.4 Quantitative PCR

Cells for RNA extraction were lysed in TRI Reagent (T3809, Merck) for 5 minutes at room temperature and RNA was extracted from TRI Reagent according to the manufacturer’s protocol. Isolated RNA was cleaned by precipitation with 3% sodium acetate in ethanol at -80°C overnight, washed as in the isolation protocol, and resuspended in H_2_O. A nanodrop spectrophotometer was used to measure RNA concentration and quality.

For cDNA synthesis, a mixture of 1μl of oligo(dT)20 (50 μM) (Invitrogen; 18418020), 2 μg total RNA, 1μl 10 mM dNTP Mix (10 mM each) (Invitrogen; 18427013), and water to make up a total of 13μl per sample was heated to 65°C for 5 minutes and incubated on ice for one minute. Next, superscript mastermix (Invitrogen; 18080093) was added to each sample (4μl 5X First-Strand Buffer, 1μl 0.1 M DTT, 1μl RNaseOUT Recombinant RNase Inhibitor (Invitrogen; 10777019), 1μl of SuperScript III RT (200 units/μl)) and the mixture was incubated at 50°C for 50 minutes and then 70°C for 15 minutes. qPCR was done in a 348 well plate, with two technical replicates per sample, and also a blank well containing no cDNA for each primer pair. Three housekeeping genes (HPRT, SDHA, RPL27) were used. Primer sequences are provided in the supplementary material. A mastermix consisting of 2μl 5x qPCR Mix Plus, 1.5μl Primer mix, and 4.5μl RNAse free per well was added to the plate. 2μl cDNA were added to each well. qPCR was run using a QuantStudio7 thermocycler with one cycle for 12 minutes at 95°C and 40 cycles of 95°C for 15s, 60-65°C for 20s and 72°C for 20s.

The data were analysed using the 2^-ΔΔCt^ method (Livak and Schmittgen, 2001). For each gene, the technical replicates were averaged. The three housekeeping genes were averaged and the ΔCt (difference between the housekeeper average and gene of interest average) was calculated for each gene of interest. The ΔΔCt was calculated as ΔCt-[Calibrator] where the calibrator is the average of the ΔCt of the controls. The final result is 2^-ΔΔCt^. This value was log-transformed prior to statistical analysis.

### 1.5 Statistical analysis

For both the ICC and qPCR experiments, three biological replicates (N=3) were analysed, where each replicate is the same cell line but with a different passage number and differentiated on a different day. The number of replicates was decided prior to the conducting of the experiments. Statistical analysis was done in Prism 9.0.2. The exposed and control mean intensity values (ICC) or log(2^-ΔΔCt^) values (qPCR) were compared using multiple 2-tailed unpaired t-tests, corrected for multiple comparisons (Holm-Šídák method). P=0.05 was used as the threshold value for statistical significance.

## 2 Results

### 2.1 Acute IFNγ exposure downregulates presynaptic genes associated with synaptic vesicles

In the RNA sequencing data from our previous study, we found downregulation of genes related to the GO term “synapses” in iPSC-derived immature neurons exposed to IFNγ for 24 hours (Warre-Cornish et al., 2020). To explore this further, a curated database of synaptic genes, SynGO (Koopmans et al., 2019), was used to identify significantly enriched biological processes (BP) and cellular component (CC) ontologies related to synaptic function. Analyses were carried out with the complete list of significantly down-regulated genes in immature neurons acutely exposed to IFNγ (25 ng/ml, 24 hours) compared with vehicle-exposed neurons. The results reveal 18 genes mapping to SynGO synaptic proteins with significant enrichment for 3 CC and 5 BP terms (Figure 1). Most of these proteins (n=12) were annotated in the presynapse cluster with 4 genes enriched for the synaptic vesicle membrane term. These results suggest that acute IFNγ exposure leads to the downregulation of 18 genes that exert presynaptic functions and regulate synaptic vesicle mechanisms in immature neurons.

**Figure 1.**
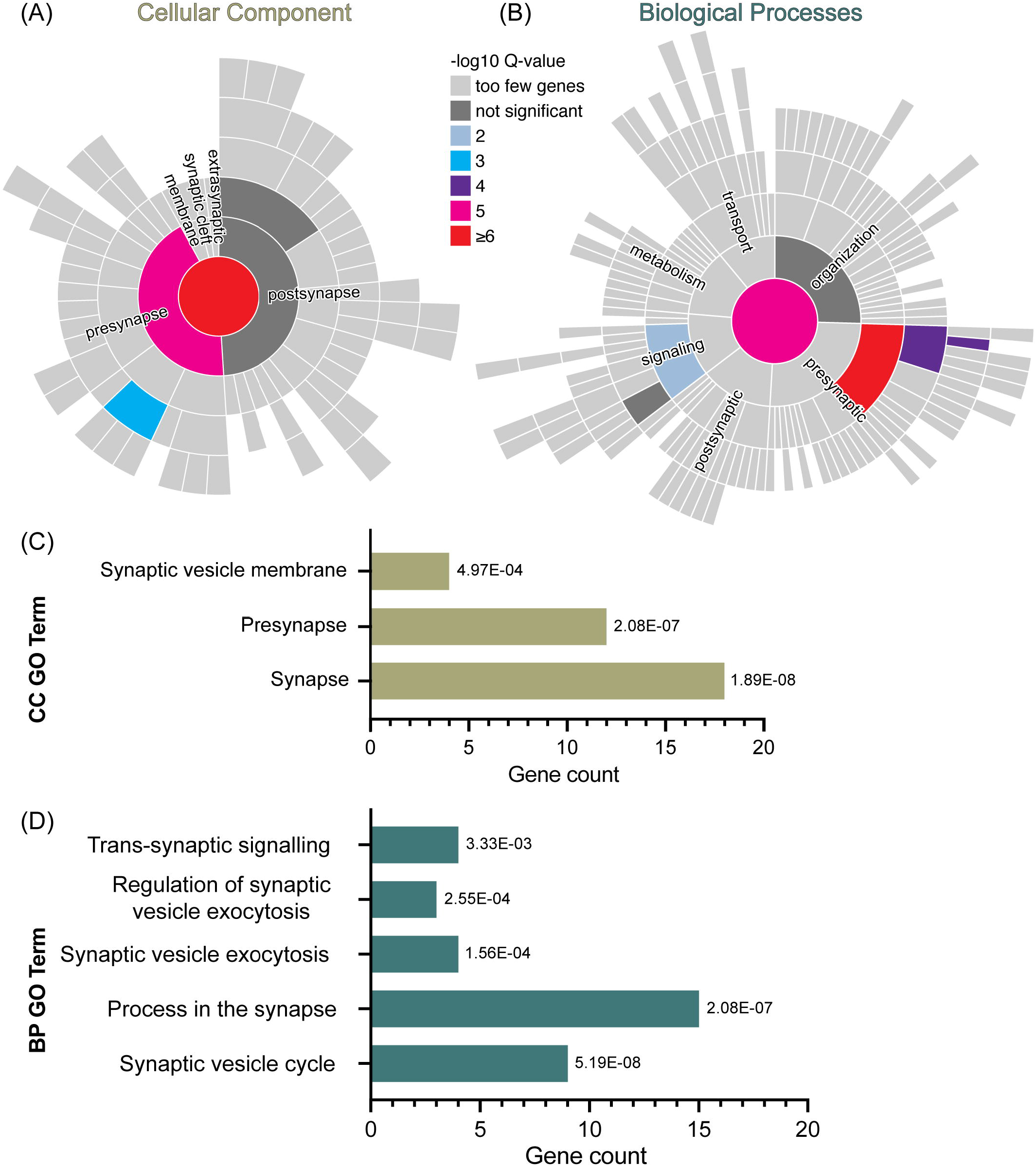
Analysis of synaptic gene ontology in immature iPSC-neurons treated with IFNγ. **(A & B)** SynGO synaptic gene ontology analyses of down-regulated DEG in D30 neurons acutely exposed to IFNγ. The sunburst plots represent synaptic annotated ontologies for CC **(B)** and BP **(A)** terms. The key colour scale indicates -log10 FDR adjusted p-values. **(A)** Significantly enriched CC ontologies include synapse (red) presynaptic (magenta) clusters. **(B)** Annotated BP terms include synaptic (magenta), presynaptic (red) and signalling (blue) terms. **(C), (D)** Plots of synaptic GO output showing the 3 CC **(C)** and 5 BC **(D)** significantly (FDR-adjusted) enriched terms for D30 IFNγ exposed neurons. The bar length indicates the number of genes, the order of each bar and numbers adjacent to each are the FDR adj. P-value. Analysis of data previously published in Warre-Cornish et al., 2020.

### 2.2 *Ngn2* overexpression generates early glutamatergic neurons at day 4

We used ioGlutamatergic line cells with *NGN2* optimized inducible overexpression to allow for rapid and reliable generation of *NGN2*-induced neurons (*NGN2*-iNs) upon treatment with doxycycline (Zhang et al., 2013;Pawlowski et al., 2017). We first validated whether the ioGlutamatergic line expresses relevant markers of glutamatergic neurons after the activation of the *NGN2* gene. By day 7 of differentiation, the cells express the pan-neuronal marker microtubule-associated protein 2 (MAP2) and excitatory presynaptic marker vesicular glutamate transporter 1 (VGLUT1) (Supplementary Figure 1). After 28 days of differentiation >99% of DAPI+ cells were immune-positive for MAP2 and also expressed TBR1, VGLUT1, CAMKIIA, and SV2A, consistent with the generation of forebrain glutamatergic neurons (Supplementary Figure 2). This is consistent with evidence that the majority of mature ioGlutamatergic neurons represent cortical excitatory neurons (Zhang et al., 2013;Lin et al., 2021). Analysis was conducted on cells at day 4 of differentiation, hereafter referred to as Day 4 *NGN2*-iNs. At this developmental timepoint, the *NGN2*-iNs generate developing immature neurons (Lin et al., 2021), consistent with the iPSC-neurons used in our previous studies (Warre-Cornish et al., 2020).

We characterized Day 4 *NGN2*-iNs using immunocytochemistry (ICC) and quantitative PCR (qPCR). We stained for the post-mitotic neuron marker neuronal nuclei antigen (NeuN) and mature neuron marker microtubule-associated protein 2 (MAP2) and the neuroprogenitor markers nestin (NES) and PAX6. In addition, staining was conducted for the immature neuron/late progenitor marker Class III β-Tubulin (TUBB3). Qualitatively, all imaged Day 4 *NGN2*-iNs expressed both the neuroprogenitor markers nestin and PAX6 and the neuronal markers NeuN and MAP2 (Supplementary Figure 3), indicating that the Day 4 *NGN2*-iNs represent early post-mitotic neurons. Morphologically, Day 4 cells had extensive processes, with some resembling developing neurons with a pyramidal cell body. Other cells had a bipolar immature morphology (Supplementary Figure 3).

qPCR for the neuronal markers *NeuN, TBR1*, and *MAP2* and the neuroprogenitor markers nestin and *PAX6* (Supplementary Figure 3) revealed that neural genes had a higher expression level compared to the progenitor genes; in particular *TBR1* and *NeuN* were highly expressed. Overall, these results indicate that Day 4 *NGN2*-iNs resemble developing immature neurons.

### 2.3 Acute exposure of Opti-OX neurons to IFN□ results in increased MHCI in the whole cell but decreased synapsin I in the cell body

Given the effects of IFN□ on synaptic genes and particularly on synaptic vesicle mechanisms, we next directly tested the effect of acute IFN□ exposure on synapsin 1, a synaptic vesicle regulator, in Day 4 *NGN2*-iNs. Synapsin I was selected as an early synaptic marker, since this protein is expressed in NPCs and colocalizes with constitutively recycling vesicles along the whole surface of developing axons that then localize to forming synapses (Zakharenko et al., 1999;Bonanomi et al., 2005).

To test how acute exposure to IFN□ affects MHCI and synapsin I expression, *NGN2*-iNs were exposed to IFN□ (25 ng/ml) or a vehicle control at day 3 for 24 hours (Figure 2A). We next asked whether these cells expressed MHCI at day 4 because MHCI has previously been used as a readout for the effect of IFN□ treatment (Warre-Cornish et al., 2020). MHCI was localised to the cell body, processes, and growth cones of all Day 4 *NGN2*-iNs (Figure 2C). All cells expressed MHCI to a similar degree. Interestingly, in all the neurons there were also bright MHCI puncta in the nucleus (Figure 2B).

**Figure 2.**
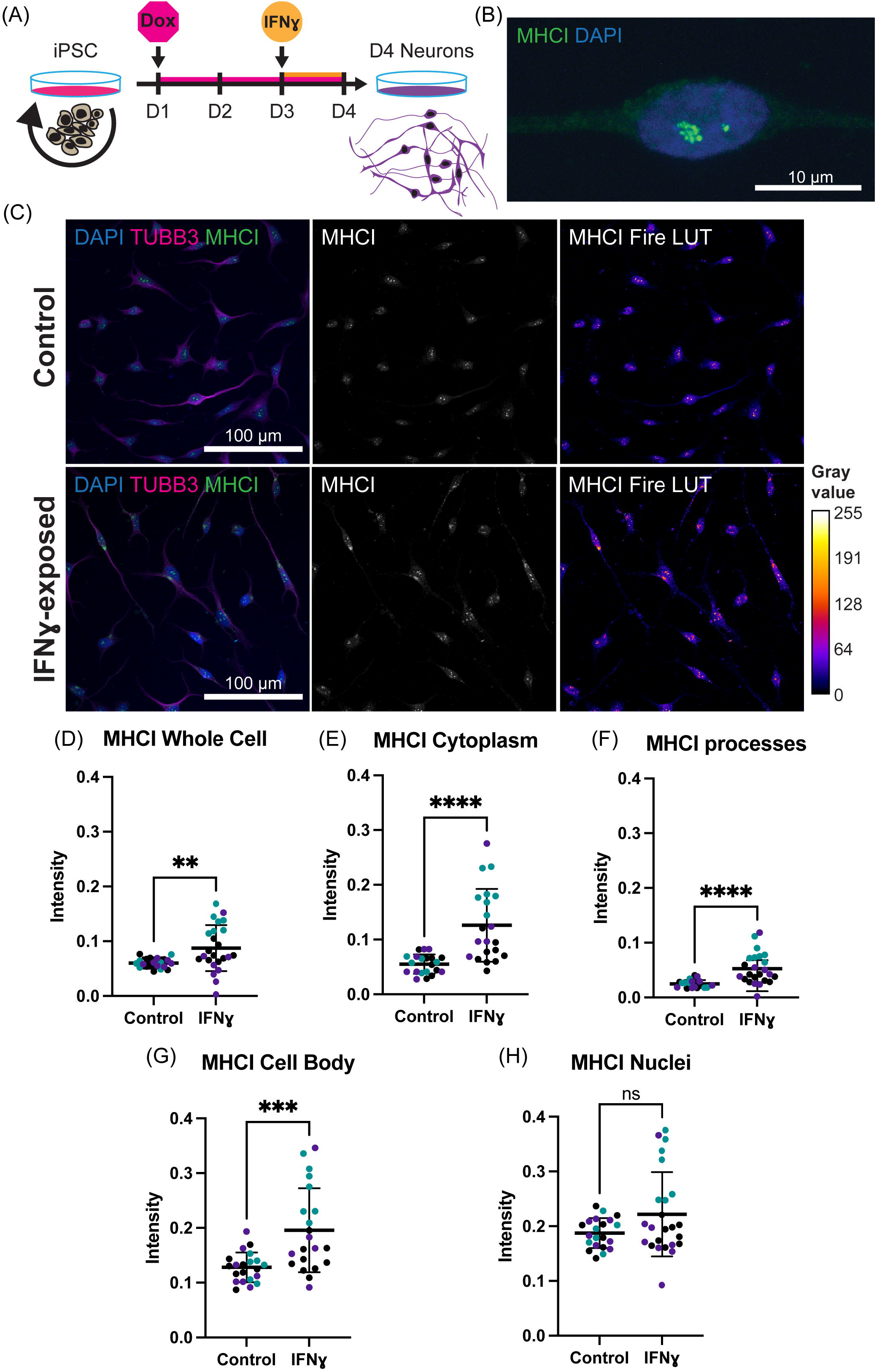
Exposure of neurons to IFN□ results in increased MHCI. **(A)** Schematic of Opti-OX neural induction and IFN□ exposure at day 3 for 24 hours. **(B)** Detailed view of a cell body, nucleus, and MHCI puncta within the nucleus. **(C)** ICC for MHCI. The top row shows control cells, the bottom row shows cells exposed to IFN□ at day 3 for 24hrs. The MHCI Fire LUT pseudo colour shows higher intensity with warmer colours and lower intensity with cooler colours. The gray values corresponding to the colours are shown on the calibration bar on the right. **(D-H)** b Scatter plots of MHCI intensity in control and IFN□-exposed neurons. The horizontal bars represent the mean, the error bars represent the standard deviation. Each point in the intensity plots represents the mean intensity of one field of view i.e., image, of the respective object. The different data point colours represent biological replicates with different passage numbers. The IFN□ and control were compared using an unpaired T-test, where N=3 and **** indicates P<0.0001, *** indicates 0.001<P>0.001, ** indicates 0.001<P>0.01, and ns indicates P≥0.05 (not significant).

Synapsin I staining was localised to the cell body, processes, and growth cones (Figure 3B). Staining was particularly bright in the cell body, with synapsin I asymmetrically localized within the shaft of one process in many neurons (Figure 3A), presumably in vesicles being transported to the processes (Figure 3A, arrowhead). Unlike MHCI, synapsin I did not localise to the nucleus and was instead at the surface of the cell within the cytoplasm. There were also sparse puncta of synapsin I along cell processes. Expression of synapsin I at day 4 is thus primarily in the cell body of all cells.

**Figure 3.**
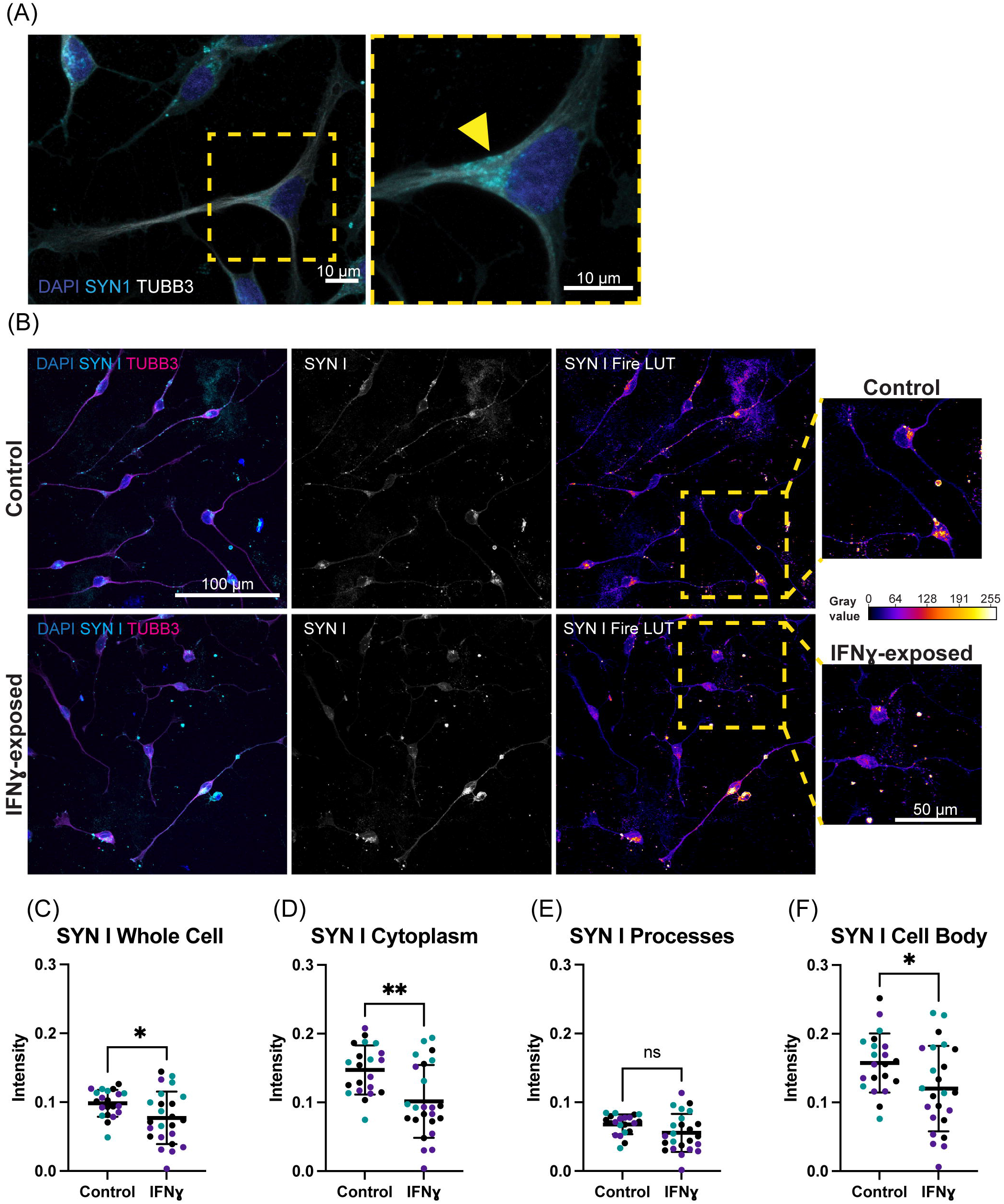
Exposure of neurons to IFN□ results in decreased SYN1 intensity in the cell bodies of some cells. **(A)** Characteristic localization of synapsin I in the cell body. Right image shows a detailed view of the highlighted region. The arrowhead indicates apparent synapsin I vesicles within the cytoplasm. **(B)** IHC for synapsin I. The top row shows control cells, the bottom row shows cells exposed to IFN□ at day 3 for 24hrs. The SYN1 Fire LUT pseudo colour shows higher intensity with warmer colours and lower intensity with cooler colours. Detailed view shown on right. The gray values corresponding to the colours are shown on the calibration bar on the right. **(C-F)** Scatter plots of synapsin I intensity in control and IFN□-exposed neurons. The horizontal bars represent the mean, the error bars represent the standard deviation. Each point in the intensity plots represents the mean intensity of one field of view i.e., image, of the respective object. The different data point colours represent biological replicates with different passage numbers. The IFN□ and control were compared using an unpaired t-test, where N=3 and ** indicates d0.001<P>0.01, * indicates 0.01<P>0.05, and ns indicates P≥0.05 (not significant).

To analyze changes in MHCI and synapsin I intensity, we designed a custom CellProfiler (Carpenter et al., 2006) pipeline to measure intensity in whole cells, nuclei, cell bodies, processes, and the cytoplasm. The IFN□-exposed Day 4 *NGN2*-iNs had a visibly higher intensity of MHCI staining compared to the control (Figure 2C). There appeared to be increased intensity of MHCI in the cell body and increased MHCI localization to the processes in the IFN□-exposed neurons (Figure 2C). Cell profiler analysis confirmed that the mean MHCI intensity was increased by 31.2% in the cells as a whole (t_(43)_=2.920, P<0.0001), increased in the cytoplasm by 56.3% (t_(40)_=4.723, P<0.0001), processes by 52.5% (t_(43)_=4.331, P<0.0001), and cell body by 34.6% (t_(40)_=3.819, P=0.0005) (Figure 2D-H). MHC I intensity in the nucleus was not significantly different (t_(43)_=1.937, P=0.06) (Figure 2H).

In contrast, synapsin I appeared to be decreased in IFN□-exposed neurons. Specifically, the asymmetrically localized clusters of synapsin I vesicles in the shaft and cell body seemed reduced in some exposed neurons, while others had intensity that is similar to control neurons (Figure 3B). Quantification showed that synapsin I intensity was decreased in the whole cell by 21.6% (t_(43)_=2.303, P=0.0261), cell body by 23.7% (t_(43)_=2.300, P=0.0263), and cytoplasm by 31.1% (t_(43)_=3.339, P=0.0017) (Figure 3C-F). The mean intensity difference in IFN□-exposed processes was not statistically significant (t_(43)_=1.840, P=0.0726, unpaired t-test) (Figure 3E). Since synapsin I does not localize to the nucleus, nuclear intensity was not analysed. These results show that IFN□ increases MHCI in Day 4 *NGN2*-iNs but has an inverse effect on synapsin I, which decreases in the cytoplasm and cell body. Cytoplasmic synapsin I and MHCI intensities in single cells are positively correlated in the vehicle condition (r=0.57, n=306), which did not change (P=0.1471, z=1.45) in the IFN□-exposed condition (r=0.49, n=389).

### 2.4 C4A and HLA-B mRNA is upregulated in IFN□-exposed neurons, but synaptic gene expression is unaltered

To validate our findings of increased MHCI staining in IFN□-exposed *NGN2*-iNs, we measured expression of *HLA-B* and *HLA-C* using qPCR. We also measured expression of the receptors for IFN□, *IFNGR1* and *IFNGR2*, and *C4A* expression. Of these, only *HLA-B* (t_(4)_=27.97, P=0.00001) and *C4A* expression (t_(4)_=11.84, P=0.000291) were significantly increased in the exposed neurons (Figure 4A).

**Figure 4.**
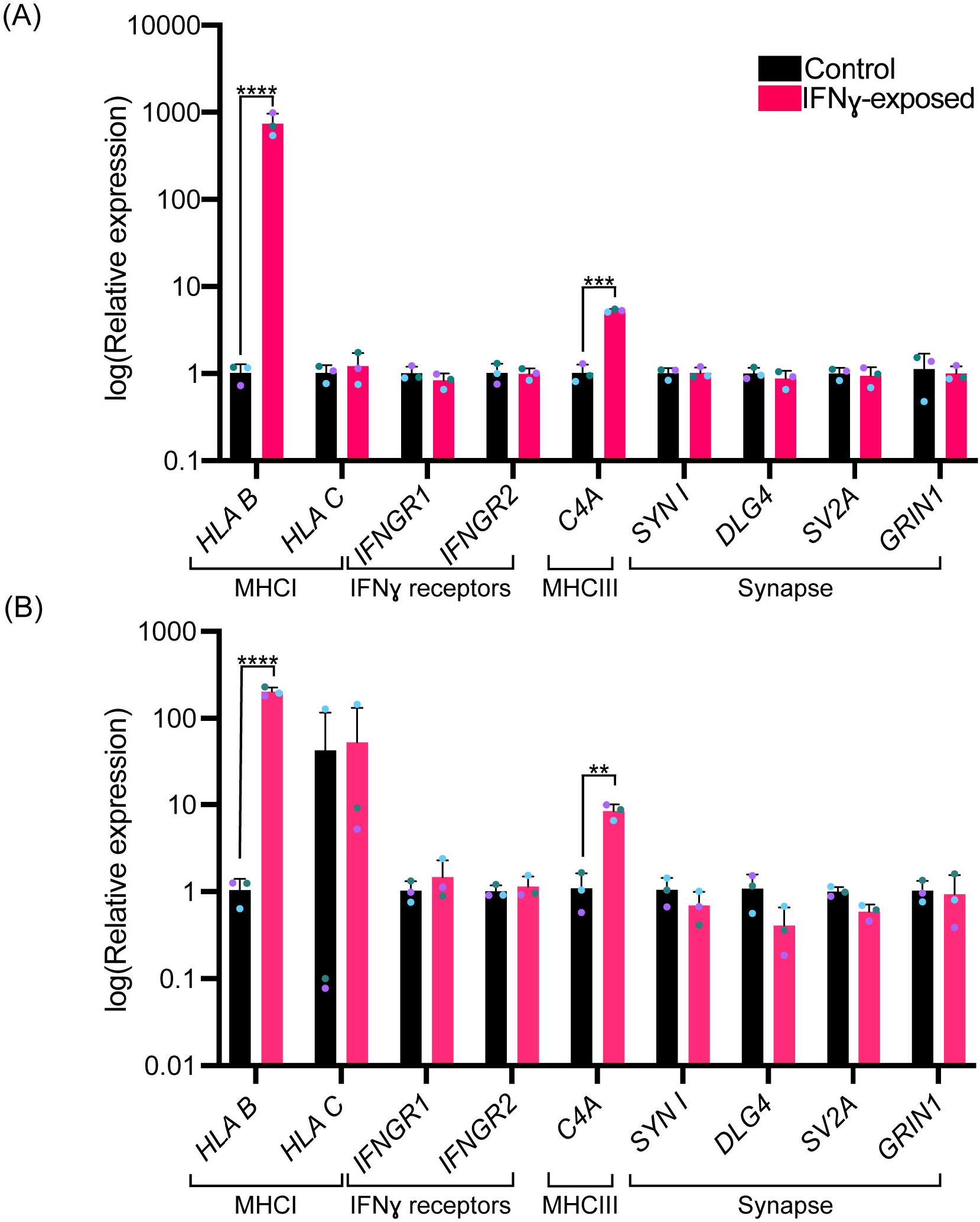
Effects of acute IFN□ treatment of gene expression in immature neurons. Bar graphs of relative expression of selected genes, showing increased *HLA-B* and *C4A* expression in IFN□-exposed neurons. The bars indicate the log(2^-ΔΔCT^), which indicates the expression relative to housekeepers and normalized to the housekeepers of the control samples (See methods for details.) **(A)** Expression in Day 4 *NGN2*-iNs exposed at day 3. N=3. **** indicates P<0.0001, *** indicates P=0.000291 (unpaired t-test). **(B)** Expression in day 27 conventionally differentiated 127_CTM iPSC line NPCs exposed at day 26. **** indicates P<0.0001, ** indicates P=0.002947(unpaired t-test). The bar represents the mean, the error bars represent the standard deviation. Points of the same colour represent the same biological replicate.

Since synapsin I decreases in the cell body following IFN□ treatment, we tested whether expression of synapsin I and other synaptic genes would be decreased following IFN□ treatment. The mean expression level for *SYN1, DLG4, SV2A*, and *GRIN1* were not significantly different (Figure 4A). We confirmed these findings using NPCs generated from a control iPSC line using a dual SMAD inhibition differentiation protocol. NPCs were exposed to IFN□ for 24 hours. *HLA-B* (t_(4)_=22.53, P=0.000023) and *C4A* expression (t_(4)_=6.466, P=0.002947) were significantly increased in the exposed NPCs (Figure 4B), which matches results from the exposed NPC-like *NGN2*-iNs.

## 3 Discussion

In this study, we used *NGN2*-iNs (ioGlutamatergic line; BitBio inc) and acute IFN□ exposure to show how synaptic protein expression can be disrupted by this cytokine. We built on studies showing IFN□ affects expression of synaptic genes in iPSC-derived immature neurons (Warre-Cornish et al., 2020).

In an acute IFN□ exposure experiment, we observed that synapsin I intensity decreased while HLA intensity increased. This is in line with published findings. For example, Glynn et al. (2011) found an increased density of clusters of synaptic vesicles with synapsin I upon siRNA knockdown of an MHCI subunit and observed significantly decreased synapsin I at inhibitory synapses when MHCI was overexpressed in rodent neurons. The decreased synapsin I we observed is thus likely linked to the concurrently increased *MHCI* expression, although we observed a positive correlation between synapsin I and MHCI intensity at the single-cell level. Decreased synapsin may translate to disruptions in synapses subsequently, as synapsin I is important for synapse maturation, including the correct localisation of synaptic vesicles in growth cones and the regulation of vesicle recycling rate (Bonanomi et al., 2005). A gene enrichment study comparing both rat (gestational day 15, LPS MIA) whole-brain and post-mortem human brain tissue samples from individuals with autism reported a common downregulation of genes associated with synaptic vesicle exocytosis (Lombardo et al., 2018). This is in line with our SynGO analysis of our IFN□ RNAseq dataset (Warre-Cornish et al., 2020) and the decrease in synapsin I associated with synaptic vesicles observed here. Whether these changes in synaptic protein translate to altered neuronal activity remains to be established. A previous study suggested that IFN□ treatment of cultured early hippocampal mouse (E15) neurons at 1-4 days *in vitro* has no effect on excitatory transmission, however it did not investigate other synapse parameters nor whether treatment of NPCs has an effect (Mirabella et al., 2021).

We did not see any difference in gene expression of our selected synaptic genes. This can be due to several possibilities. There may be changes in synapsin I translation, mRNA turnover, expression of splice variants, or misregulation of genes associated with synapsin I. Expressed protein reflects an earlier time point than mRNA expression, the absence of any difference in mRNA levels could therefore also indicate a transient change in expression that is no longer detectable after 24 hours. The synapsin I intensity change observed may also reflect changes in the spatial distribution of synaptic proteins or in local translation, which is not detectable in whole-cell RNA.

The increase in MHCI expression and staining intensity following IFN□ exposure matches the findings of a previous neuroprogenitor cell study that used the same 24 hour acute IFN□ exposure of iPSC-NPCs and -neurons (Warre-Cornish et al., 2020). In this study, Warre-Cornish et al. (2020) found persistent upregulation of MHCI gene expression and increased MHCI protein levels. They also described upregulation of *HLA-C* and *HLA-B* expression; however, we only observed a significant increase in *HLA-B*. As in the study, we observed no change in the expression of IFN□ receptors. MHCI is known to be involved in synaptic plasticity and learning (Datwani et al., 2009;Shatz, 2009;Lee et al., 2014;Adelson et al., 2016) and is important for negatively regulating synapses (McAllister, 2014). Dysregulation of *MHCI* expression could thus potentially be sufficient for a downstream disruption of synapses even if no change in synaptic genes is present at the point of IFN□ exposure. MHCI has been shown to mediate reduced synaptic connectivity in a mouse MIA model by signalling through myocyte enhancer factor 2 (MEF2) (Elmer et al., 2013). Future work would need to establish whether these changes in MHCI expression persist and whether other downstream changes arise as the neurons mature.

We also observed increased expression of the complement component *C4A* after acute IFN□ exposure, in line with Warre-Cornish et al. (2020). *C4A* mRNA levels are increased in post-mortem schizophrenia patient brains and *C4A* variants are associated with schizophrenia risk (Sekar et al., 2016). Genes downregulated upon increased C4A expression are enriched for schizophrenia risk (Kim et al., 2021). C4A is expressed by neurons and colocalizes with synaptic markers and is thought to mediate pruning of synapses (Sekar et al., 2016). Overexpression of C4A in mice resulted in behavioural changes, reduced cortical synapse density, and increased engulfment of synapses by microglia (Yilmaz et al., 2021). Inhibition of microglial activity reverses MIA abnormalities, including synapse loss (Ikezu et al., 2021). Increased *C4A* expression could therefore contribute to decreased synapse number. Co-culture studies with microglia would be particularly informative for future IFN□ exposure studies.

The ioGlutamatergic cell line provides a good system for future IFN□ exposure studies of neurons. Our findings suggest a possible link between IFN□ exposure and reduced synaptic vesicles in early neurons, potentially via MHCI, however more work is needed to understand the basis of this link.

## 5 Author Contributions

DPS and ACV: conception and design, literature searching, manuscript writing and editing and project supervision. AP, RM, LS, LDP and NA: carried out experiments. AP: manuscript writing and editing. DPS and ACV: financial support. All authors approved the final manuscript.

## 7 Acknowledgments

The authors thank Daniel Beglin for his insightful comments on image analysis. The authors also thank the Wohl Cellular Imaging Centre (WCIC) at the IoPPN, Kings College, London, for help with microscopy.

