## Supplemental Material for "Interferon-□ Exposure of Human iPSC-derived Neurons Alters Major Histocompatibility Complex I and Synapsin I Protein Expression"

#### *Supplementary Material*

- **Supplementary Table 1**
- **Supplementary Figures 1-3**

### 1 Supplementary Figures and Tables

List of primers for qPCR and their sequences

| Gene Name | Forward Primer (5'-3') | Reverse Primer (5'-3') |
|---|---|---|
| SYN1 | CTGGACGTCCCAAACCAC | CTTTCACCTCGTCCTGGCTA |
| HLA B | TGAGATGGGAGCCGTCTT | ACCTGAACTCTTCCTCCTACA |
| HLA C | GCTGACCGAGTGAGCCT | TGTTCTTCTTTGATAGCCCATGA |
| DLG4 | GTGACGACCCATCCATTTTC | CTGCCTCTTTGAGGGCTTC |
| SV-2A | GACGGTGTGGAGGTCTTTGT | CGAAGACGCTGTTGACTGAG |
| GRIN1 | ACTCGGACAAGAGCATCCAC | TCCAGCTGTAGACACGCATC |
| IFNgR1 | GGTCTGTGAAGAGCCGTTGTC | CGGGACCACGTCAGGAATAT |
| IFNgR2 | GGAAAAGGAGCAAGAAGATGTTCT | AGCTCCGATGGCTTGATCTC |
| C4A | GTTGCTCTTGTTCTCTCCTTCT | CACTGATCCTTTCACTACCTGTC |
| HPRT | TGACACTGGCAAAACAATGCA | GGTCCTTTTCACCAGCAAGCT |
| SDHA | AGGAATCAATGCTGCTCTCTGG | CTGCTCCGTCATGTAGTGGA |
| RPL27 | ATCGCCAAGAGATCAAAGATAA | TCTGAAGACATCCTTATTGACG |
| TBR1 | ACACTGGGGCTCACTGGAT | AAAACCACCATCTGCCCATT |
| PAX6 | GCCAGAGCCAGCATGCAGAACA | CCTGCAGAATTCGGGAAATGTCG |
| MAP2 | CTCTCGCACAGAGTTATCC | GACCTACCACCAAGTCCTAAAC |
| TUBB3 | CAAGATGTCCTCCACCTTCATC | GACACCAGGTCGTTCATGTT |
| RBFOX3 | CCTTCCACCGTTTCCTTCTC | GACTTGGTTGGATGCCTCTTATC |
| Nestin | GAAACAGCCATAGAGGGCAAA | TGGTTTTCCAGAGTCTTCAGTGA |

#### 1.1 Supplementary Figures

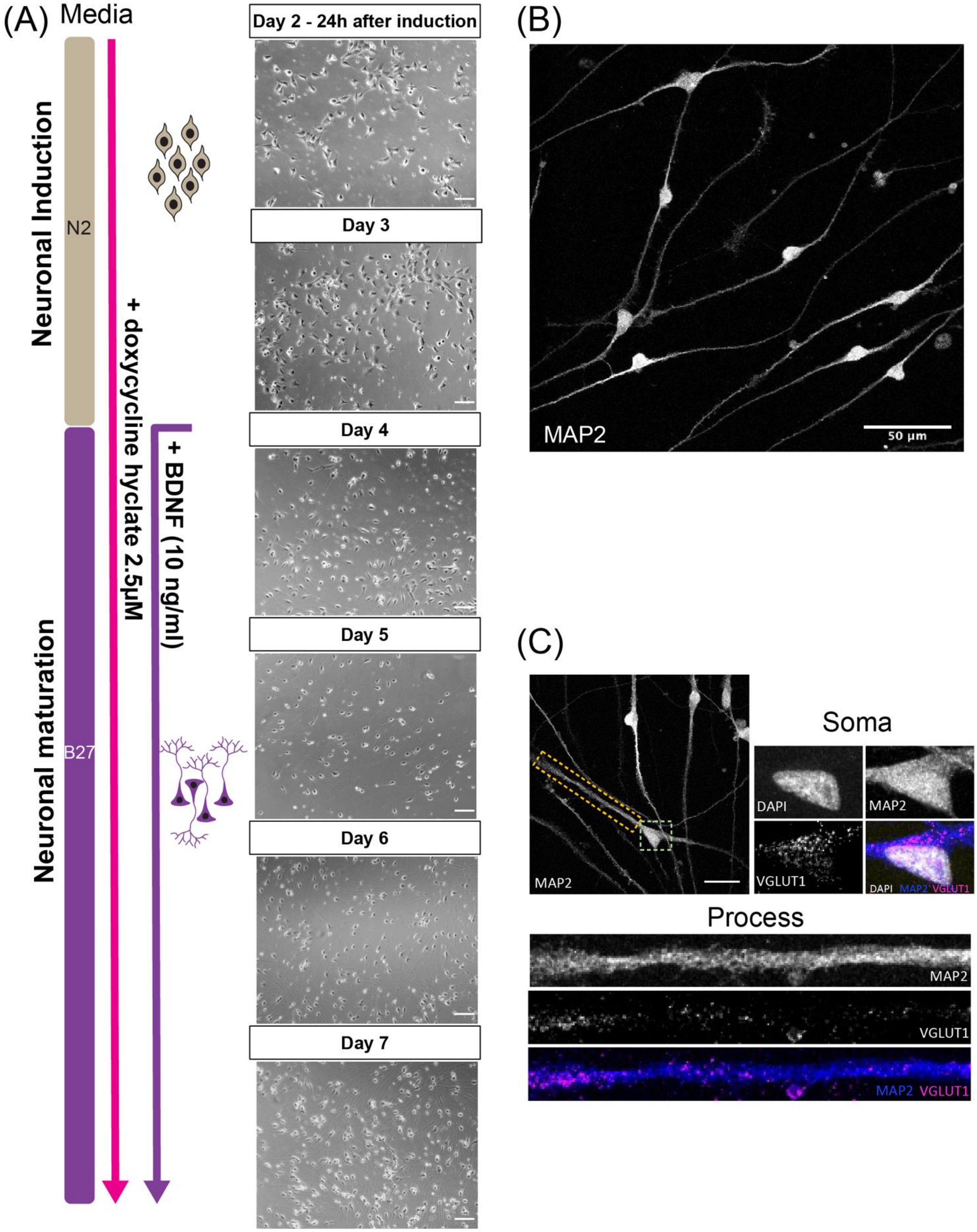

**Supplementary Figure 1. ioGlutamatergic line rapidly generates glutamatergic neurons following neural induction. (A)** Schematic representation and timeline of differentiation protocol. Brightfield images were acquired using an EVOS XL core imaging system with a 10x objective. The scale bars represent 150 µm. **(B)** Representative confocal image of neurons on day 7 after neuronal induction immunostained for neuronal marker microtubule-associated protein 2 (MAP2). Image was acquired using a Leica SP5 confocal microscope with a 63x oil-immersion objective. **(C)** Representative confocal image of neurons on day 7 after neuronal induction immunostained for neuronal marker microtubule-associated protein 2 (MAP2; in blue) and excitatory presynaptic marker vesicular glutamate transporter 1 (VGLUT1; in magenta). The scale bar represents 25 µm. Dotted lines indicate zoomed in regions showing soma and process. At this stage of early neuronal maturation, VGlut1 localises to somatic and process regions. Image was acquired using a Leica SP5 confocal microscope with a 100x oil-immersion objective.

(A)

(B)

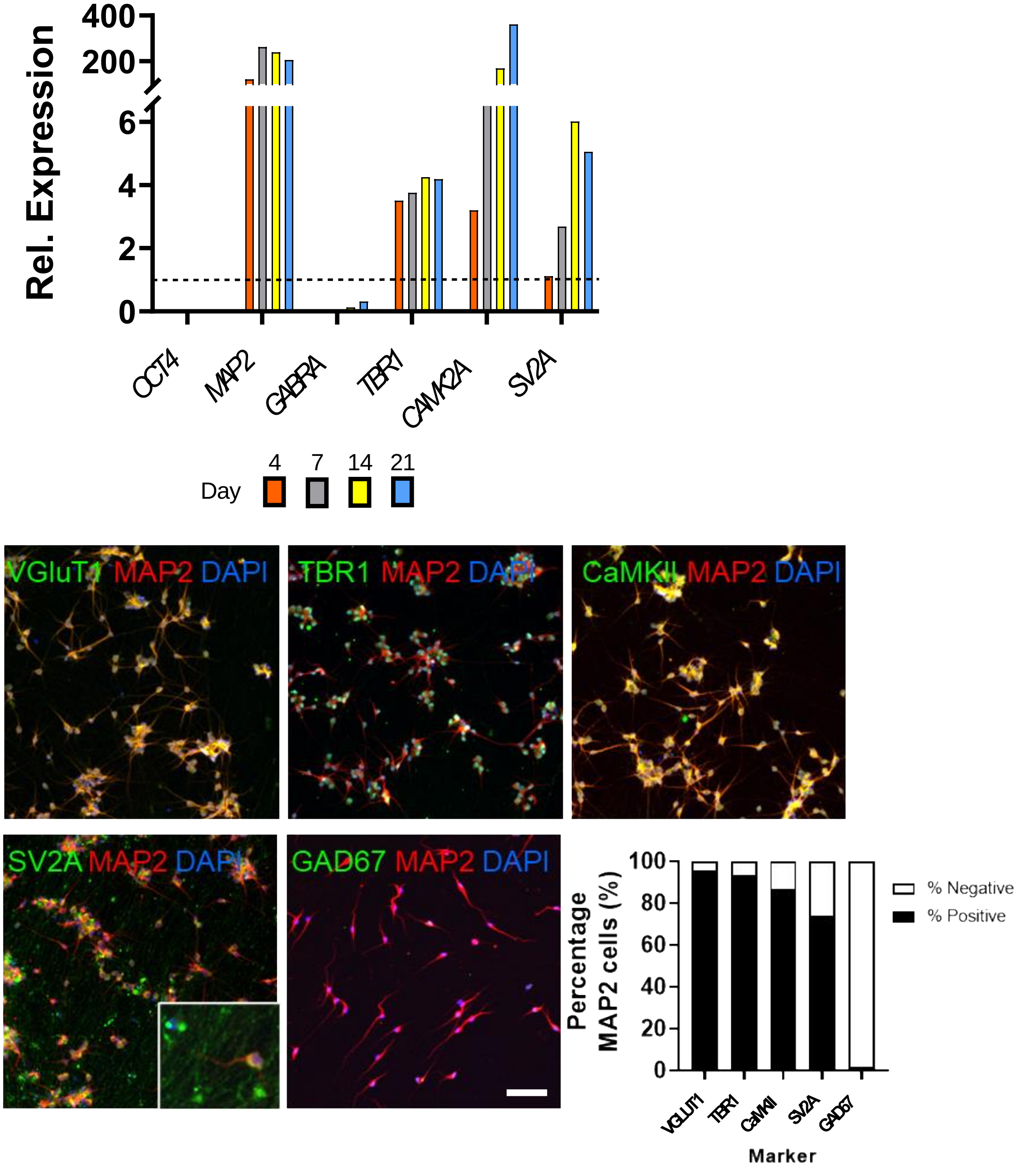


**Supplementary Figure 2. Generation of glutamatergic neurons from ioGlutamatergic line. (A)** Time-dependent increase in mRNA expression of key fate and synaptic genes indicating the development of excitatory neuronal properties. All expression levels have been normalized to the day 0 time point as a relative expression of 1, indicated by the dotted line.

**(B)** After 28 days of differentiation >99% of cells are positive for MAP2 and expressed TBR1, VGLUT1, CaMKII and SV2A consistent with the generation of glutamatergic neurons. In addition, neurons were found to be negative for the inhibitory neuronal marker, GAD67. Representative images acquired using Thermo Scientific CellInsight High-Content Screening Platform. The scale bar represents 200µm.

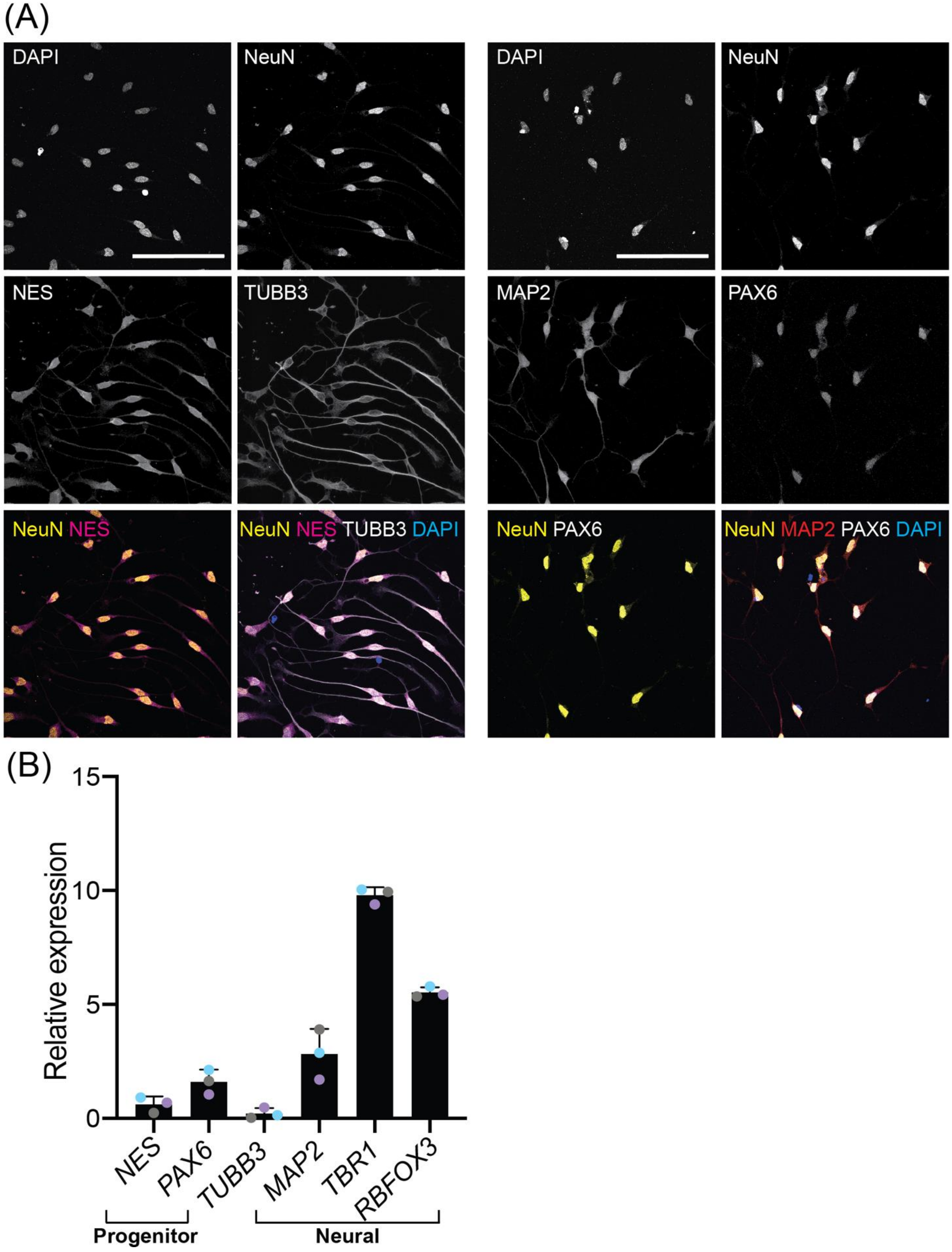
(A)
DAPI
NeuN
DAPI
NeuN
NES
TUBB3
MAP2
PAX6
NeuN NES
NeuN NES TUBB3 DAPI
NeuN PAX6
NeuN MAP2 PAX6 DAPI
(B)
Relative expression
15
10
5
0
NES
PAX6
TUBB3
MAP2
TBR1
RBFOX3
Progenitor
Neural

**Supplementary Figure 3. Expression levels of neural and neuroprogenitor genes in D4 ioGlutamatergic cells**. **(A)** NeuN, nestin (NES), MAP2, PAX6, and Class III β-Tubulin (TUBB3) expression in day 4 *NGN2*-iNs. Representative images acquired using a Leica SP5 confocal microscope with a 63x oil-immersion objective. The scale bar represents 100µm. **(B)** Bar graph of the relative expression of neural and neuroprogenitor genes, the expression level is the ΔCt value relative to housekeepers. The bar represents the mean, the error bars represent the standard deviation. Points of the same colour represent the same biological replicate.
